## Supplementary materials for "Repeatability of adaptation in sunflowers: genomic regions harbouring inversions also drive adaptation in species lacking an inversion"

### Supplementary Methods:

*Whole genome re-sequence data alignment and variant calling*

DNA was extracted from leaf tissues (these are the same individuals examined in the previous study, see Todesco et al. (2020) for details). Briefly, all libraries were sequenced at the Genome Quebéc Innovation Center on HiSeq2500, HiSeq4000 and HiSeqX instruments, to produce paired end, 150 bp reads (Illumina, San Diego, CA, USA). Libraries with a low number of reads were re-sequenced to increase genome coverage. After quality filtering (see below), a total of 60.7 billion read pairs were obtained. Illumina adapters and poor quality reads were hard-clipped using Trimmomatic (v0.36) (Bolger et al. 2014). Reads were then aligned to the *H. annuus* XRQv1 genome (HanXRQr1.0-20151230; Badouin et al. (2017)) using NextGenMap (v0.5.3; (Sedlazeck et al. 2013)). PCR duplicates were marked and removed using (picard MarkDuplicates 2.9.3). Genomic regions containing transposable elements (~3/4 of the sunflower genome) were excluded to reduce computational time and improve variant quality. Genotyping for each species was performed independently, as joint-genotyping on the whole ensemble of samples was computational impractical. GATK's VariantRecalibrator (v4.0.1.2; Van der Auwera and O'Connor 2020), which filters variants in the call set according to a machine learning model inferred from a small set of "true" variants, was used to remove low-quality calls and produce a dataset of a more manageable size. In the absence of an externally-validated set of known sunflower variants to use as calibration, we computed a stringently-filtered set from top-N samples with highest sequencing coverage for each species (N=67 for cultivated sunflower, and N=20 for wild sunflower species). The stringency of the algorithm in classifying true/false variants was adjusted by comparing variant sets produced for different parameter values (tranche 100.0, 99.0, 90.0, 70.0, and 50.0). For each cohort, results for tranche = 90.0 were chosen for downstream analysis, based on heuristics: the number of novel SNPs identified, and improvements to the transition/transversion ratio (towards GATK's default target of 2.15).

*Remapping sites to the HA412-HO reference genome*

As described with details by Todesco et al. 2020, haploblock analysis highlighted contig ordering issues with the XRQv1 reference assembly (see below). To overcome this, all sites were transferred to a new reference, HA412-HOv2, which used Hi-C for contig and scaffold ordering (Belton et al. 2012; Marie-Nelly et al. 2014). To do this, the 200 bp of reference sequence flanking each site in XRQv1 were extracted and aligned to HA412-HOv2 using BWA (Li 2013). These alignments were filtered for mapping quality > 40 and the HA412-HOv2 position for the variant site was extracted. Since all remapped sites were not in repetitive regions and had passed VQSR filtering, remapping success rate was high (96-98%). Whenever mapping suggested two different variants on the XRQv1 genome were in the same position on the HA412-HOv2 genome, likely due indels and imprecise alignment, one site was shifted by one bp so they did not overlap. Remapping was preferred to *de novo* read alignment and variant calling against the HA412-HOv2 assembly because of the prohibitive amount of computational time that would have required.

*Gene ontology enrichment analysis of regions with repeated association*

Genes that overlapped with CRAs associated with environmental and phenotypic variables were screened for enrichment of Gene Ontology (GO) terms. GO annotations for *Arabidopsis thaliana* genes from the TAIR database were mapped onto their sunflower homologs and a custom database of sunflower GO annotations was constructed. The R package TOPGO (Alexa and Rahnenfuhrer 2022) was used to analyze the set of candidate genes to determine which categories were most overrepresented. Significance for each individual GO identifier was computed with Fisher's exact test and significant GO terms were identified at an FDR of 1%. GO functional enrichment analysis was performed in the categories biological process (BP), cellular component (CC) and molecular function (MF).

### Supplementary Results

**GO-enrichment analysis.** To investigate the functional associations of genes overlapping with the windows of repeated association (WRA), we performed gene ontology (GO) analysis using TopGO package in R. GO terms with corrected P-values < 0.05 were considered significantly enriched. Supplementary Figure S8 & S9 provide a list of GO terms that are over-represented in our gene set. GO components and processes associated with membrane assembly, transport through the endomembrane system, mRNA and growth are significantly enriched in this analysis.

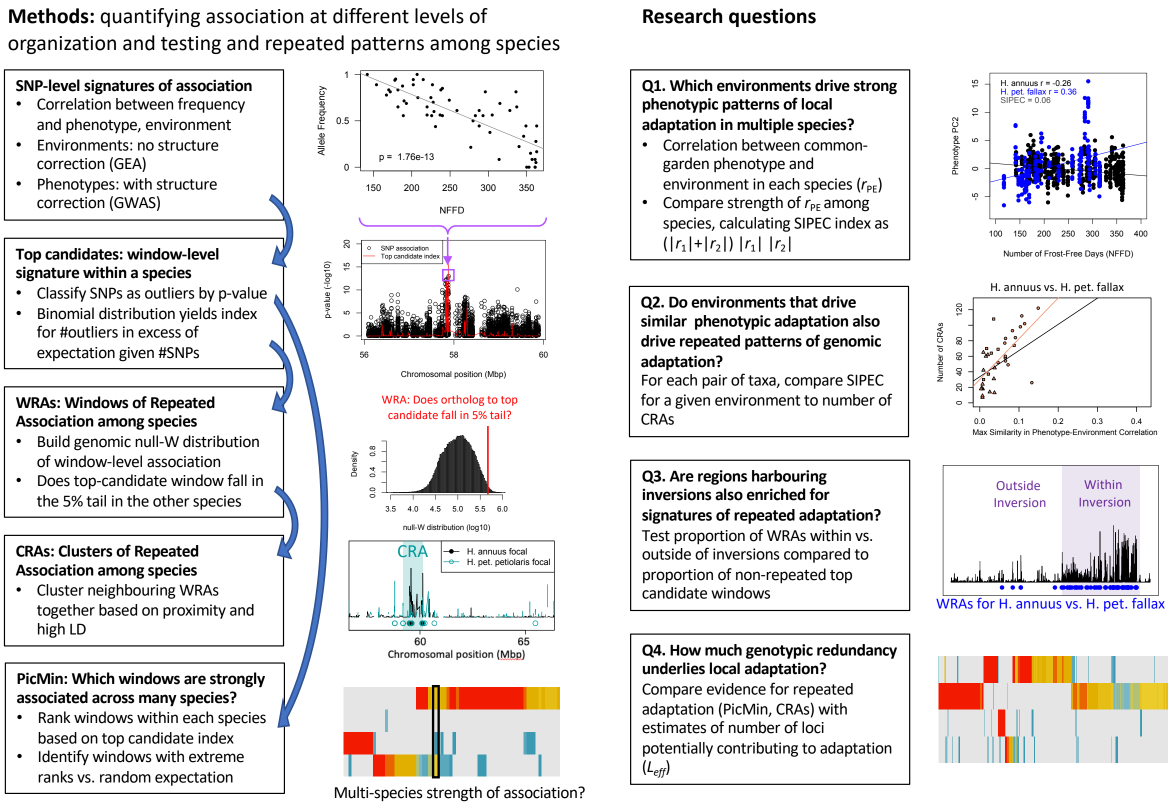


**Supplementary Figure S1.** Schematic overview of methods and primary research questions

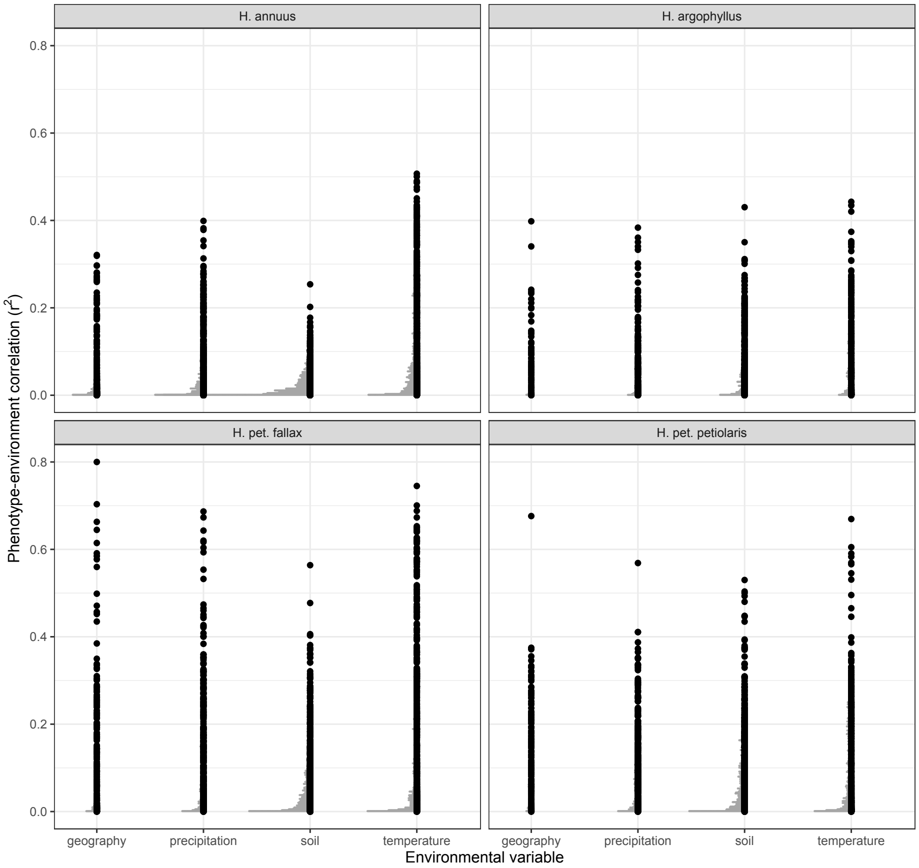


**Supplementary Figure S2.** Strength of phenotype-environment correlations across all traits for four different types of environmental variables, in each of the sunflower species and subspecies. Black points show individual values, grey points show binned density.

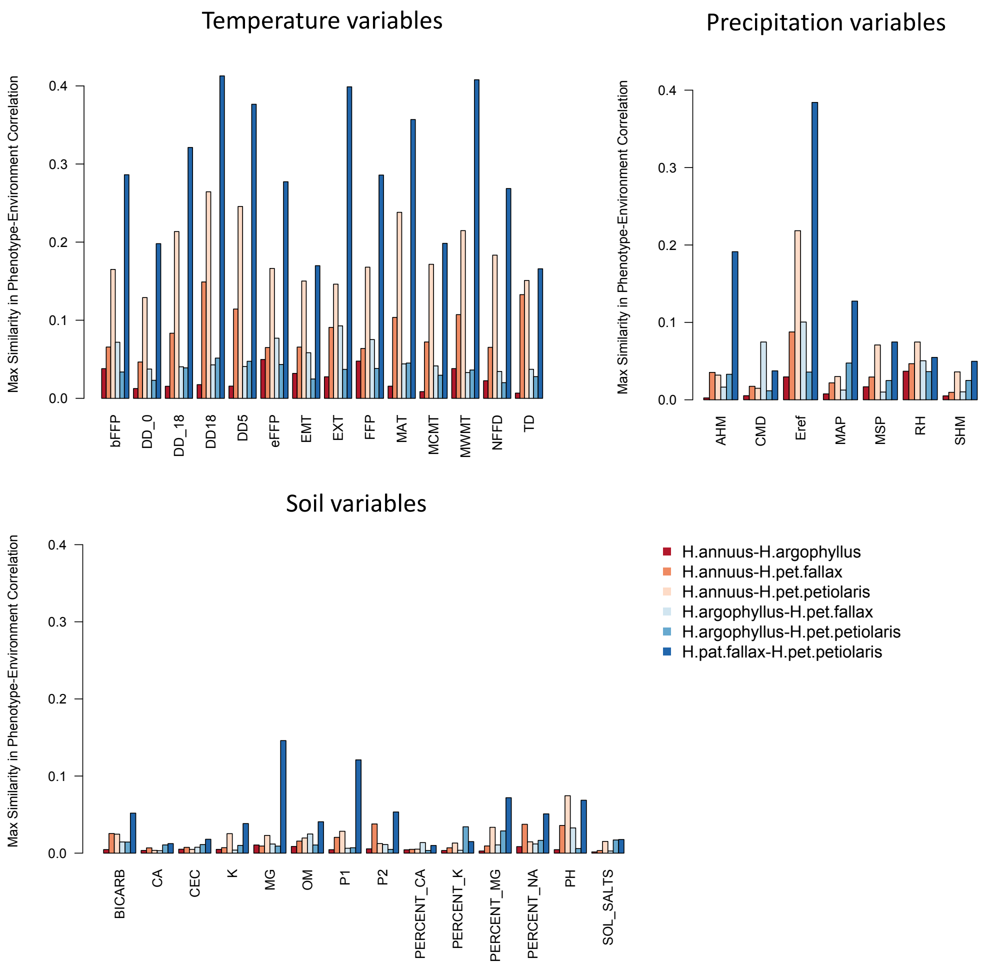


**Supplementary Figure S3.** Maximum Similarity in Phenotype-Environment Correlation (SIPEC) for pairs of taxa, across soil-, temperature-, and precipitation-related environmental variables. The maximum value of SIPEC for each environment is calculated across all phenotypic PCA axes that cumulatively explain 95% of the variance.

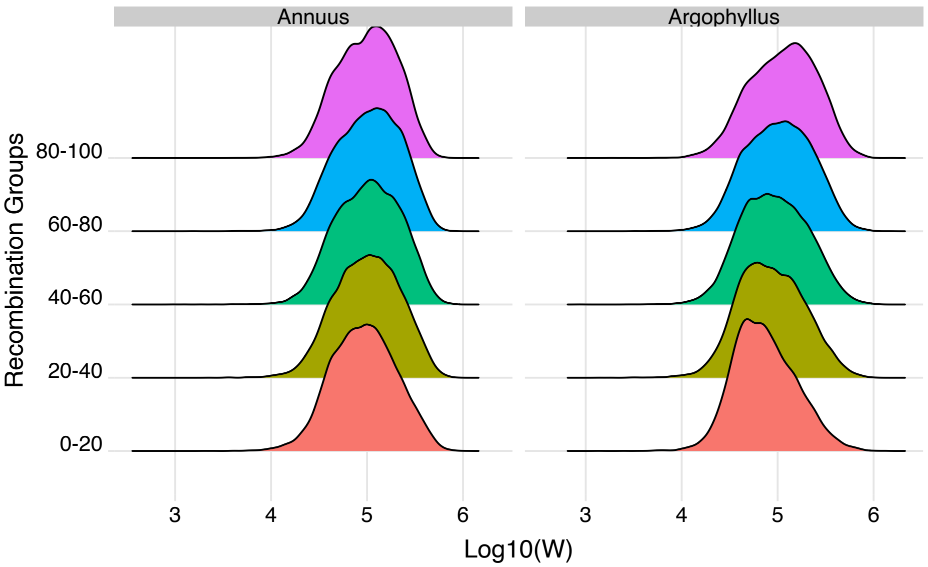


**Supplementary Figure S4.** The effect of recombination rate on width of the null-W distribution for the NFFD variable for *Helianthus annuus* and *H. argophyllus*. Recombination bins represent the $0^{th}$-$20^{th}$ percentile, $20^{th}$-$40^{th}$ percentile, etc.

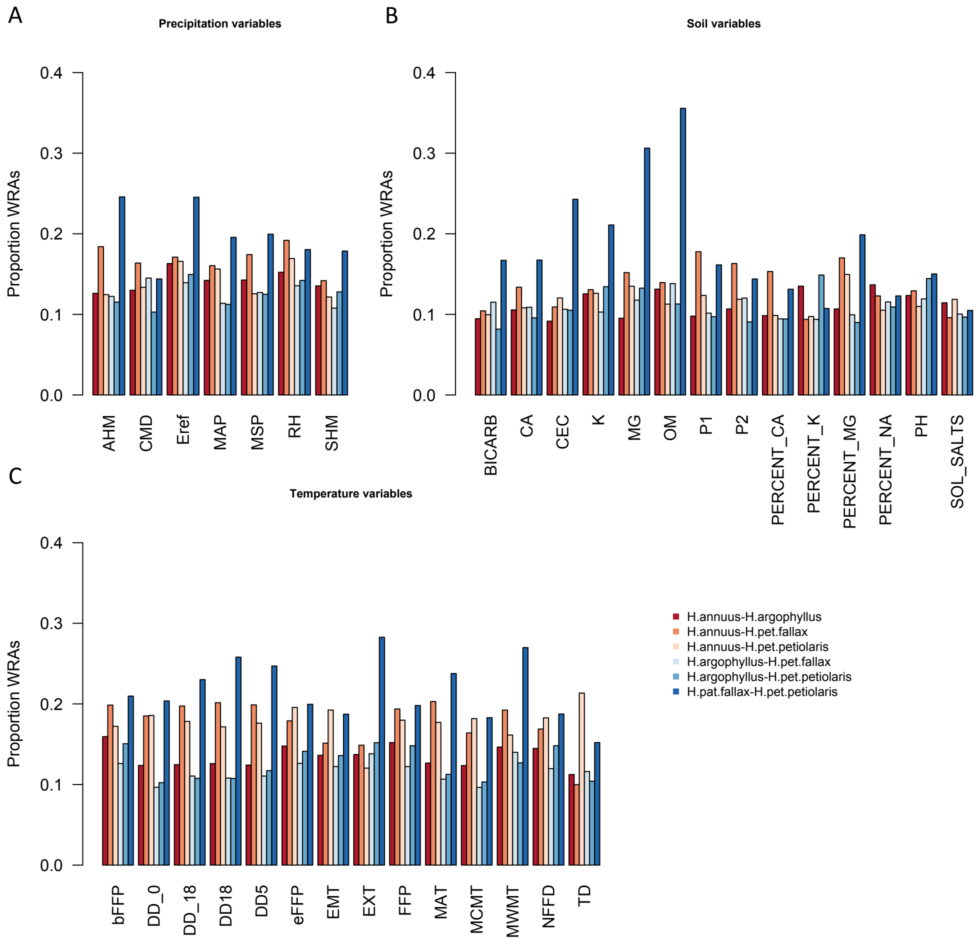


**Supplementary Figure S5.** Proportion of top candidate windows that are significant hits under the null-W test (Windows of Repeated Association), for pairs of taxa, across geographic and soil-, temperature-, and precipitation-related environmental variables.

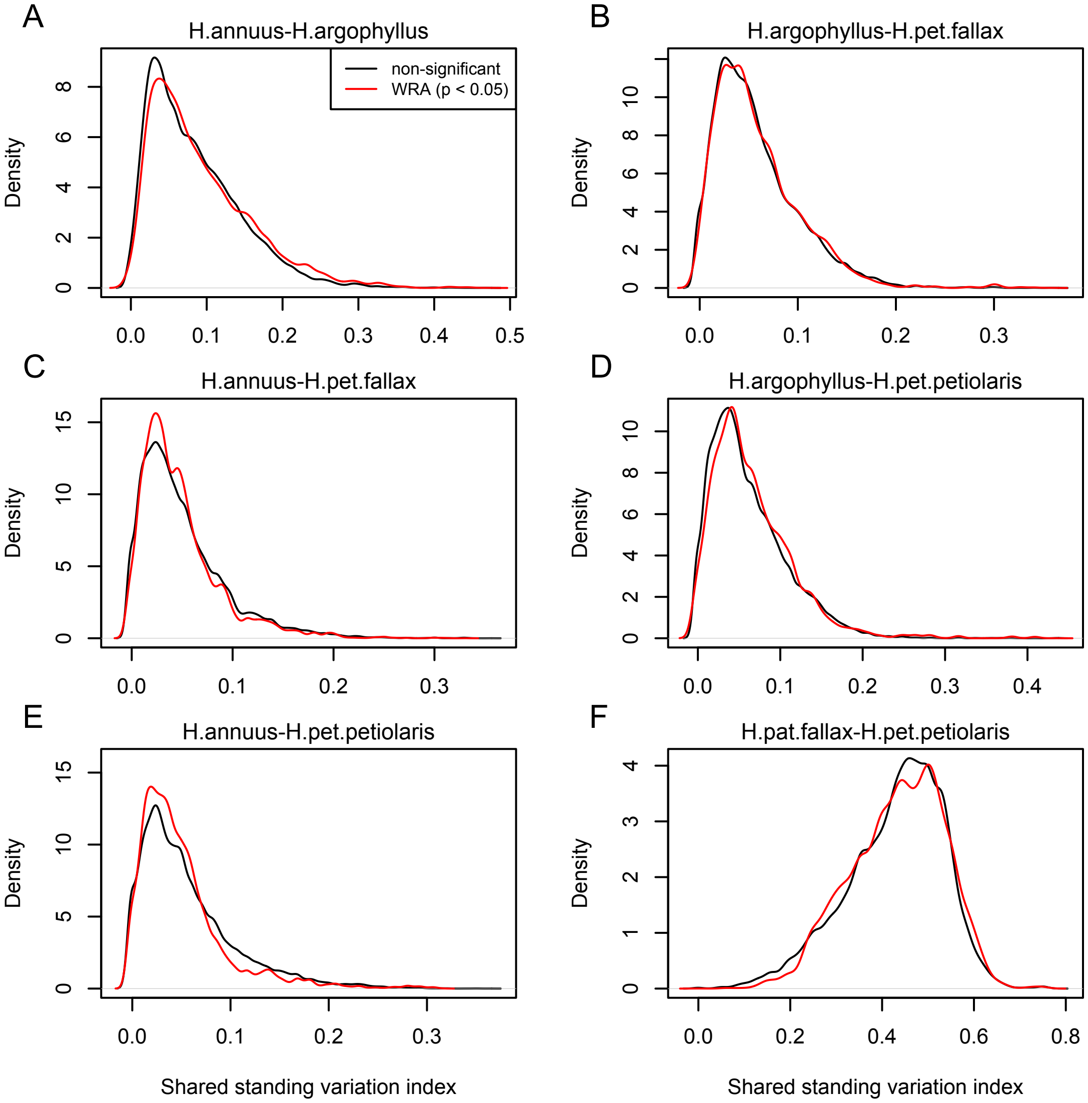


**Supplementary Figure S6.** Index of shared standing variation for Windows of Repeated Association (WRAs) vs. top candidates that were not significant under the null-W test. The index of shared standing variation reflects the proportion of SNPs that are shared vs. non-shared among species and provides an indicator of the likely extent of introgression, which does not appear to differ substantially among the windows of the genome with significant ($p < 0.05$; red lines) vs. non-significant ($p > 0.05$; black lines) null-W test results.

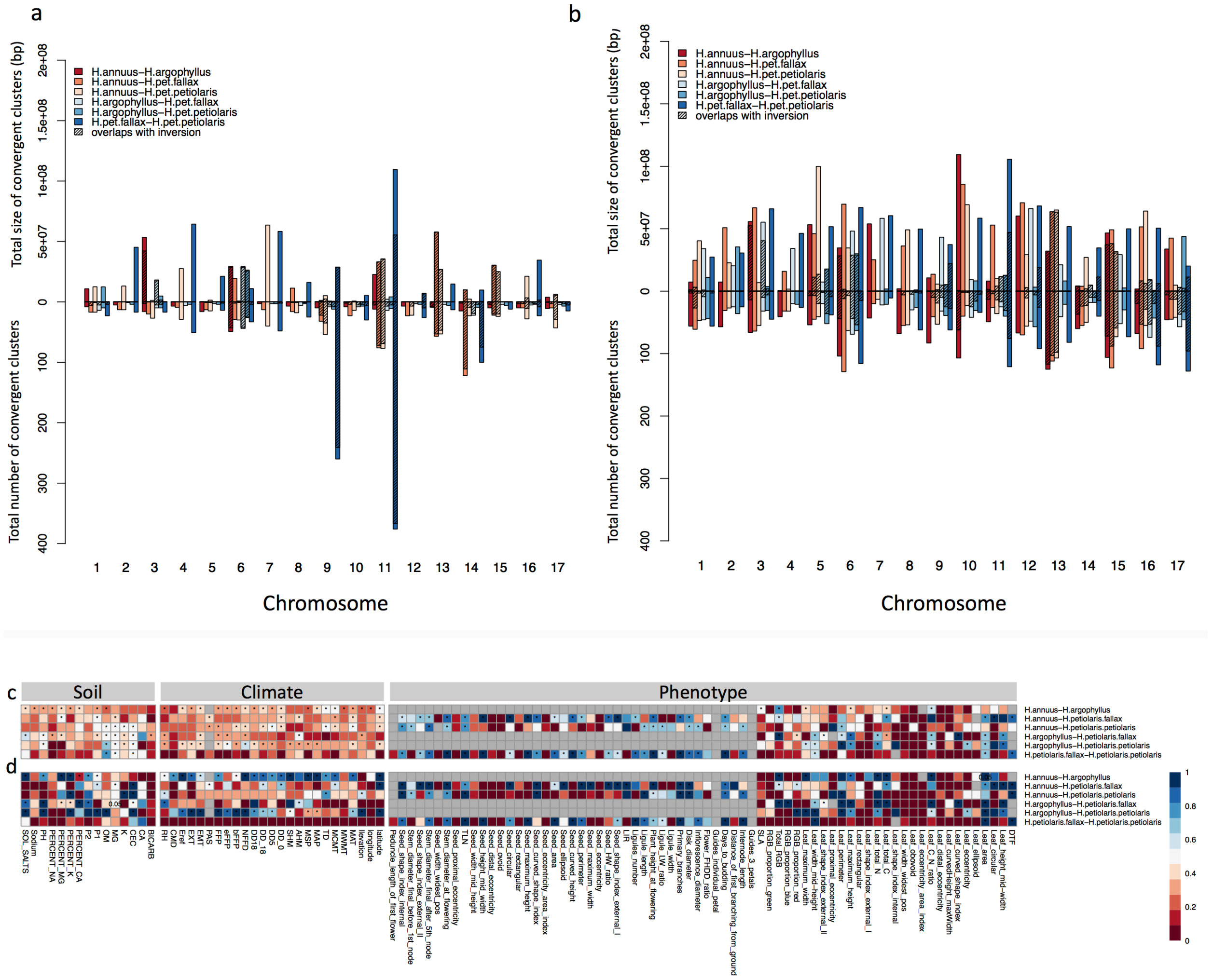

**Supplementary Figure S7.** Number and size of Clusters of Repeated Association (CRAs) and their overlap with haploblocks. Total size and total number of CRAs detected among six studied pairs on each linkage group across all phenotypes by GPA (a), and environmental GEA (b). Hatching areas indicate the total size and number of clusters residing within chromosomal rearrangements. Heat maps present proportion of CRAs by number (c) and size (d) per each phenotype variable and environment variable overlapping with chromosomal rearrangements. Stars in indicate overlaps between CRAs and haploblocks happen significantly different from chance (P-value <= 0.05). Gray cells in the heat maps indicate no data is available for that comparison and variable.

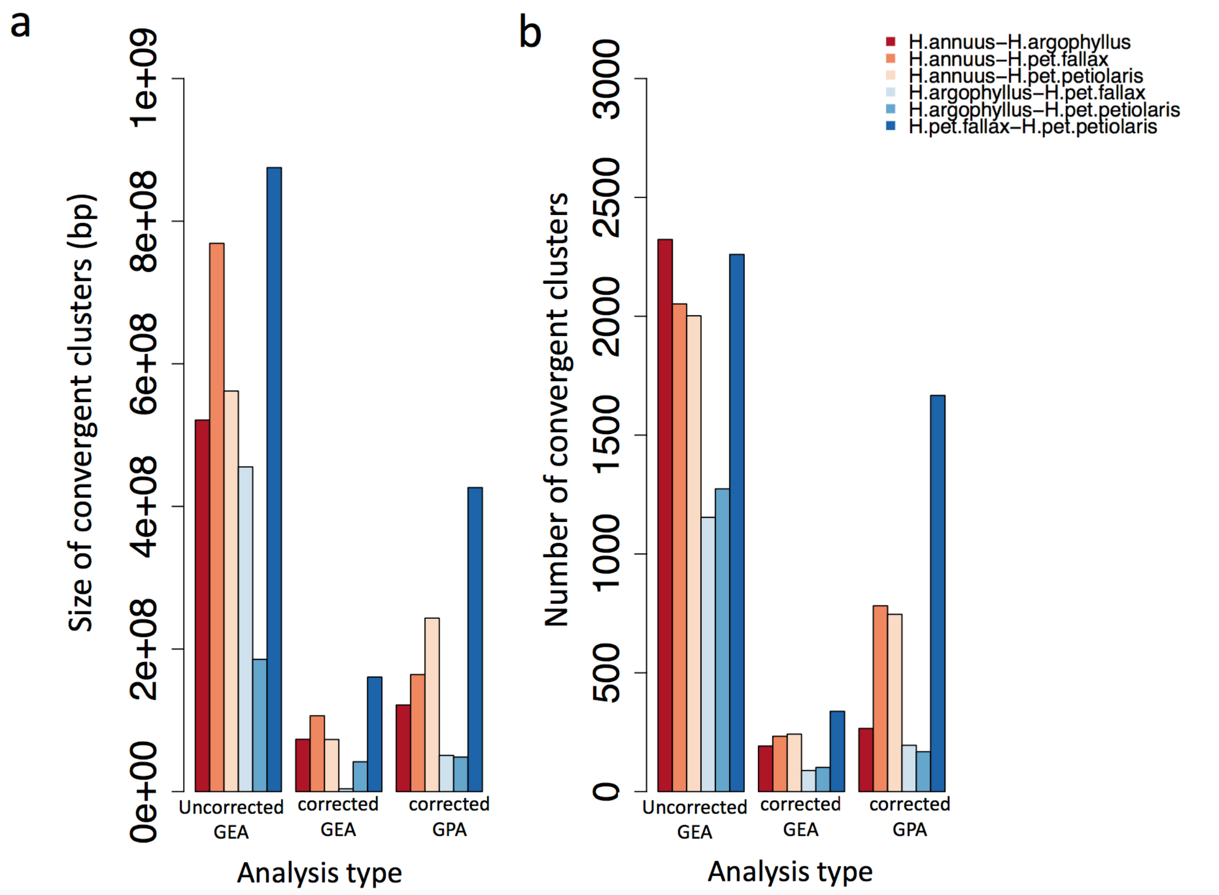


**Supplementary Figure S8.** Total size and number of convergent clusters in different pairs for each analysis type. The total size of convergent clusters (a) and total number of convergent clusters (b) identified among different pairs surveyed in the present study using association genetic approaches that corrected population structure versus those that did not correct across all environmental variables (GEA) and corrected GPA.

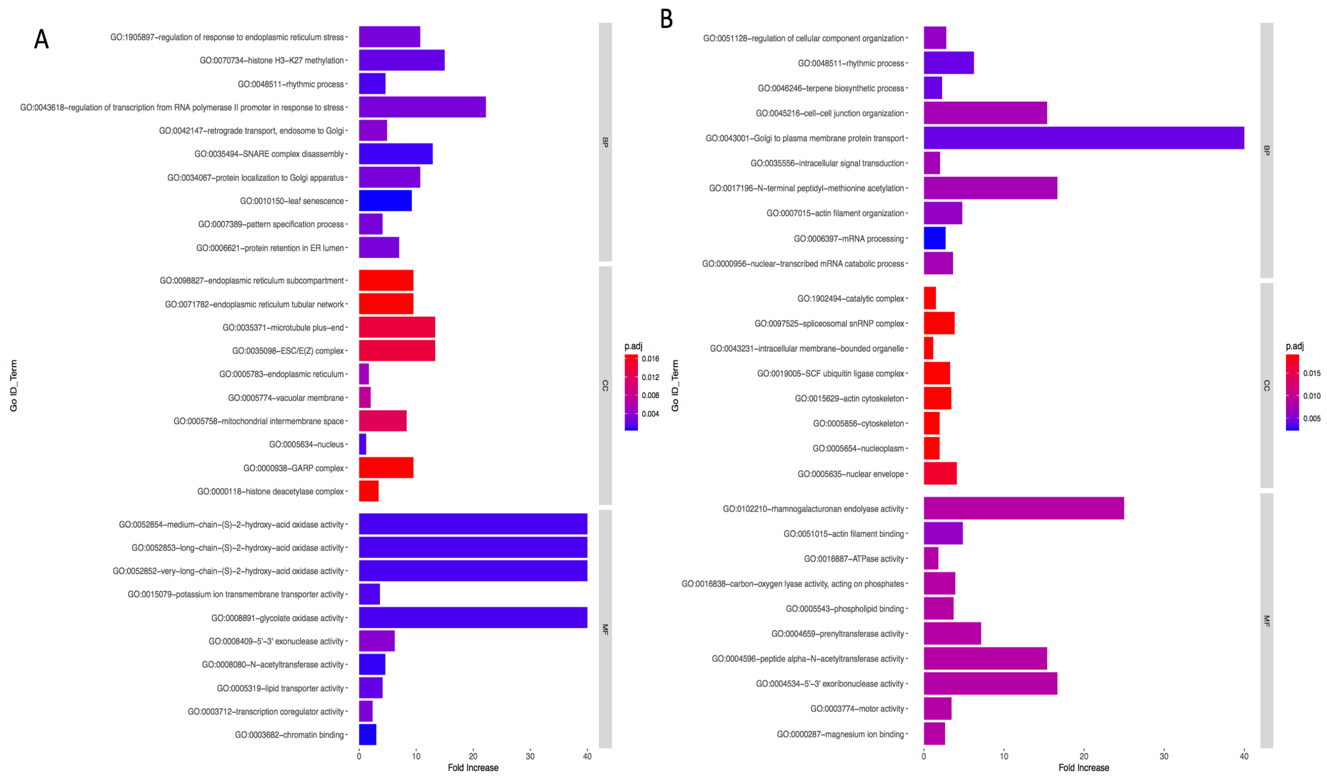


**Supplementary Figure S9.** Bar graph of Gene Ontology (GO) enrichment analysis for phenotype (A) and precipitation related variables (B). Bar plot depicts the significant enriched gene ontology (GO) terms within categories: biological process, cellular component, molecular function. Y-axis represents the GO term, and the X-axis represents the enrichment significance, respectively

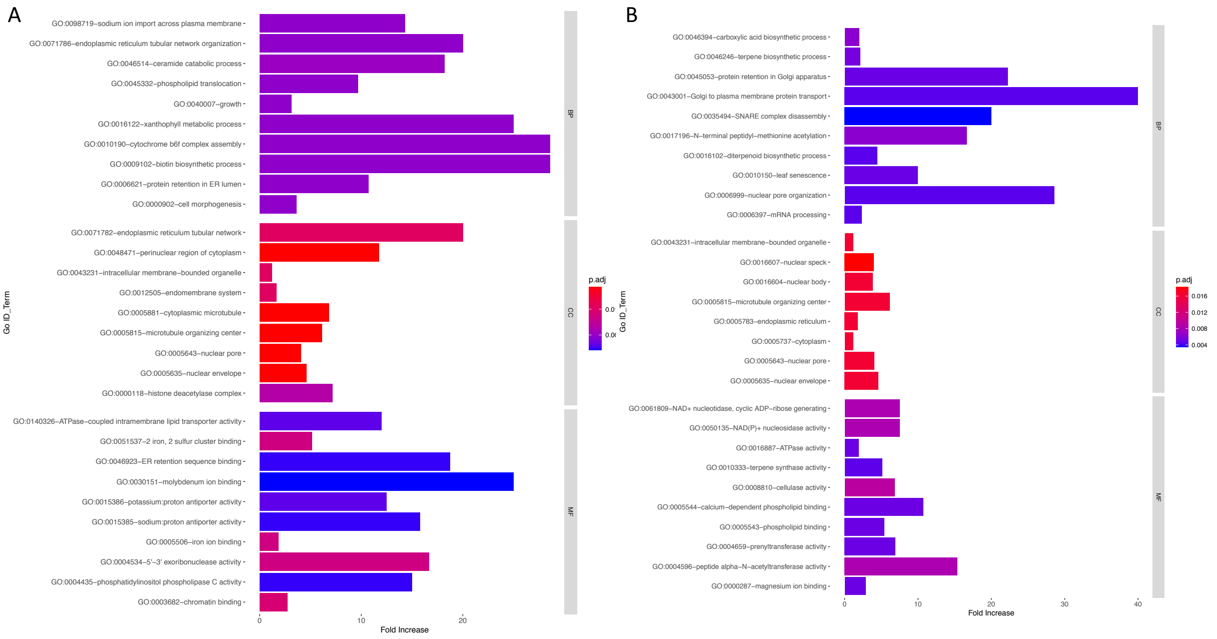


**Supplementary Figure S10.** Bar graph of Gene Ontology (GO) enrichment analysis for soil (A) and temperature related variables (B). Bar plot depicts the significant enriched gene ontology (GO) terms within categories: biological process, cellular component, molecular function. Y-axis represents the GO term, and the X-axis represents the enrichment significance.

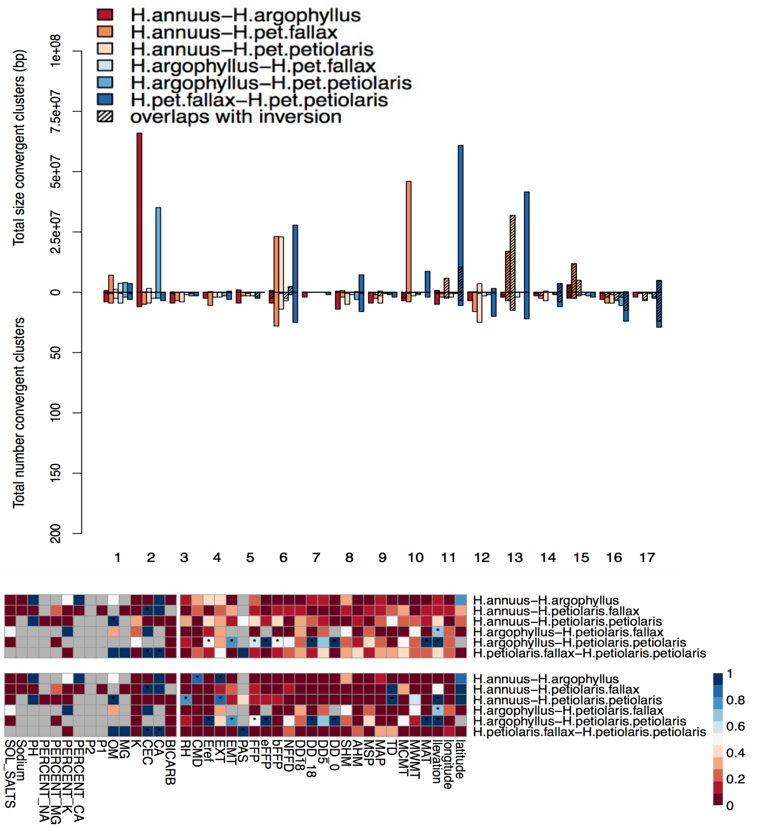


**Supplementary Figure 11.** Effect of structure correction on number and size of Clusters of Repeated Association (CRAs) and their overlaps with inversions. Total size and total number of CRAs detected among six studied pairs on each linkage group across all environmental variables by corrected GEA (a). Hatching areas indicate the total size and number of clusters residing within haploblocks. Heat maps present proportion of CRAs overlapping with haploblocks by number (b) and size (c) for each phenotype and environmental variable (climate and soil). Stars indicate cases where observed overlaps between CRAs and haploblocks happen significantly more than expected by chance (P-value <= 0.05). Gray cells in the heat maps indicate no data is available for that comparison and variable.

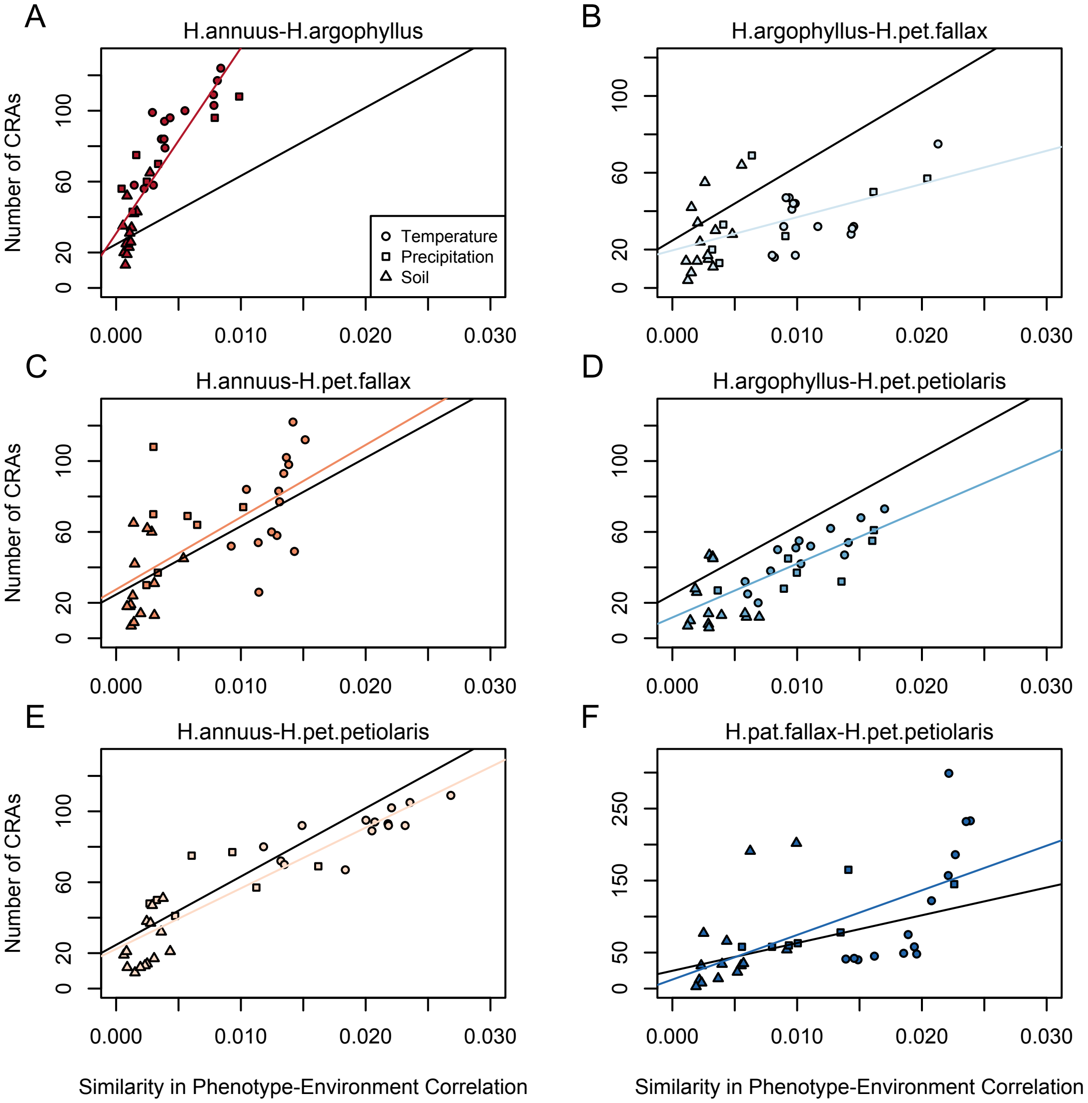


**Supplementary Figure S12.** Relationship between mean Similarity in Phenotype-Environment Correlation (SIPEC) and number of Clusters of Repeated Association (CRAs). SIPEC was calculated for each phenotypic PCA axis with the mean calculated across the axes that cumulatively explain 95% of the phenotypic variance, for each environment. Each panel includes both a linear model fit to the data within the panel (coloured lines), and a linear model fit to all data simultaneously (black lines) for comparison. Note that because environmental variables are correlated, these points are not independent and therefore represent a source of pseudoreplication, preventing formal statistical tests of this relationship.

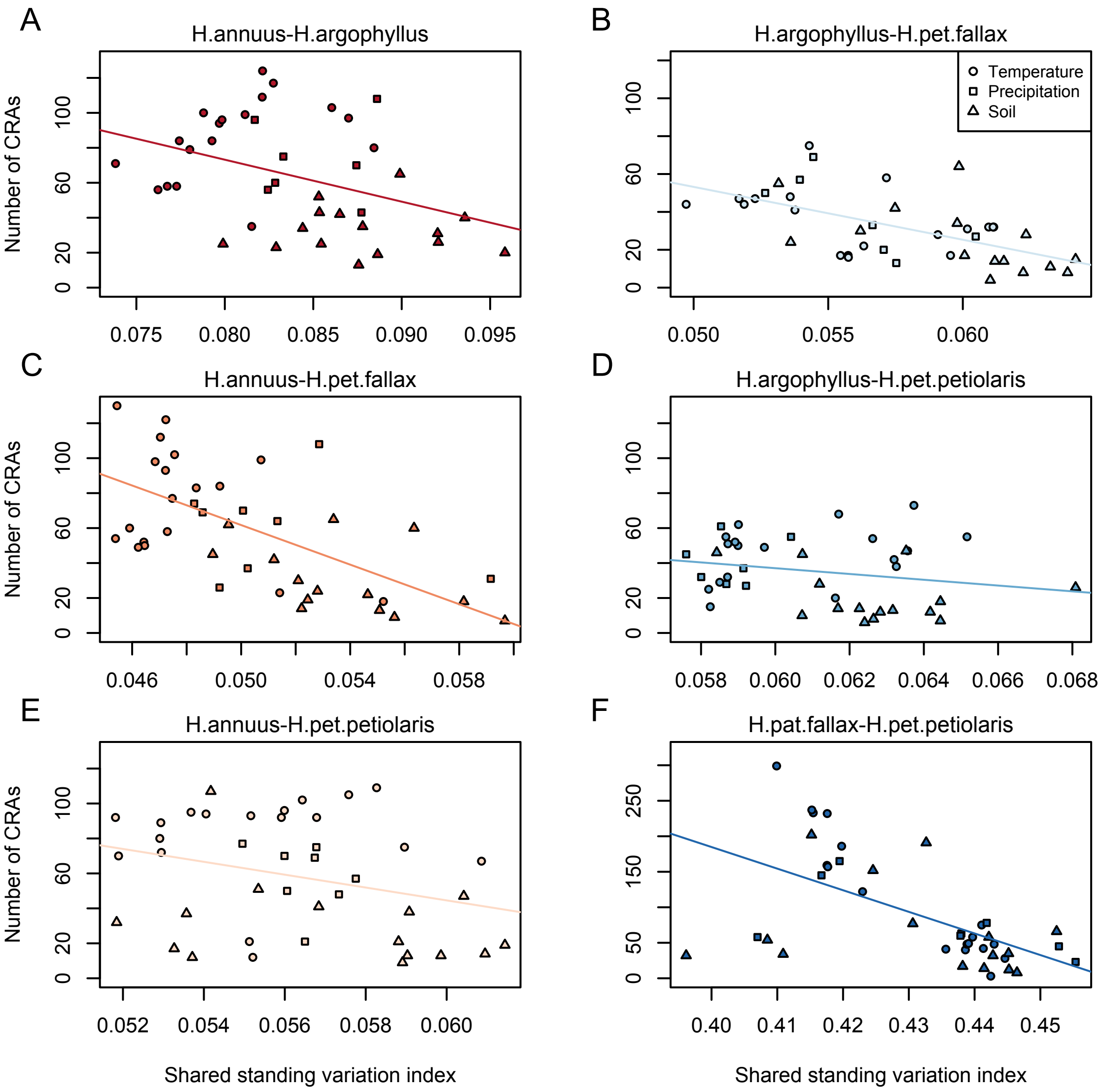


**Supplementary Figure S13.** Relationship between index of shared standing variation and number of CRAs. Lines show linear model fits for data within each panel.

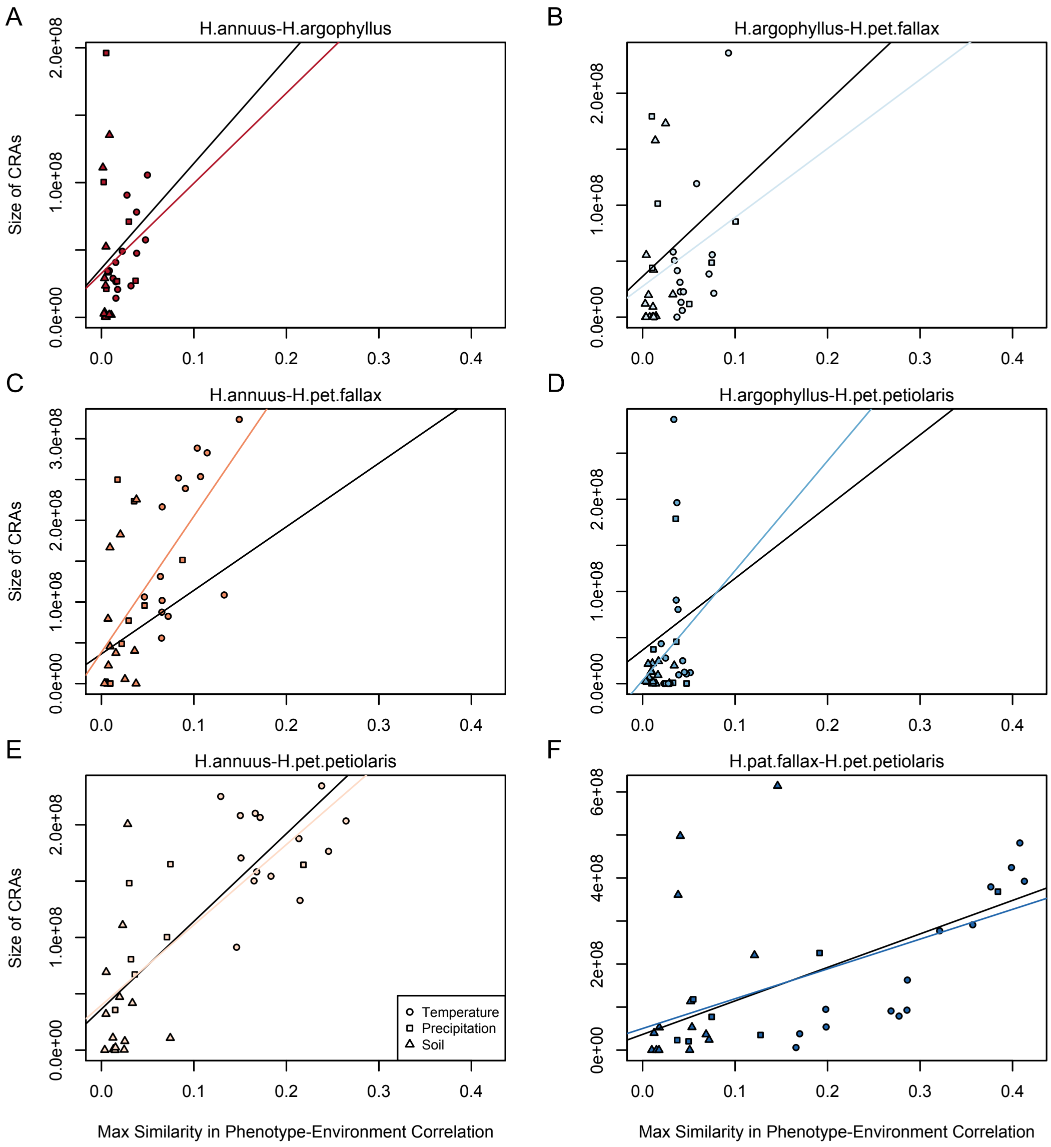


**Supplementary Figure S14.** Relationship between mean similarity in phenotype-environment correlation (SIPEC) and size of Clusters of Repeated Association (CRAs). SIPEC was calculated for each phenotypic PCA axis with the mean calculated across the axes that cumulatively explain 95% of the phenotypic variance, for each environment. Each panel includes both a linear model fit to the data within the panel (coloured lines), and a linear model fit to all data simultaneously (black lines) for comparison. Note that because environmental variables are correlated, these points are not independent and therefore represent a source of pseudoreplication, preventing formal statistical tests of this relationship.

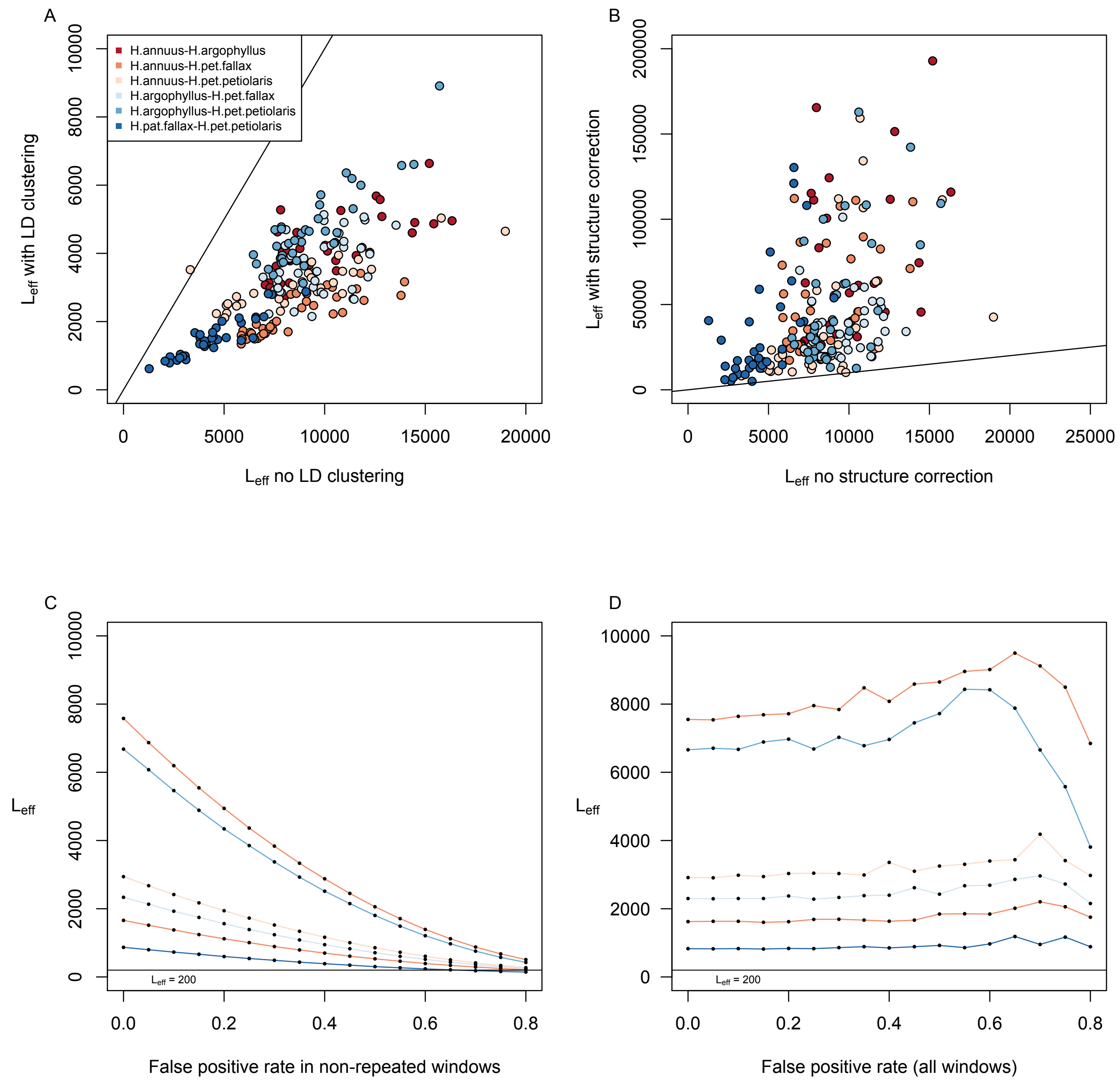


**Supplementary Figure S15.** Estimates of effective number of loci in pairwise contrasts among species. Panel A shows a comparison of estimates of the effective number of loci ($L_{eff}$) when calculated with vs. without LD-clustering for the environmental variables from the 6 pairwise contrasts among lineages. Panel B shows the effect of structure correction using Baypass on $L_{eff}$. Panels C & D show the estimation of $L_{eff}$ for the variable with the lowest average value (Hargreaves reference evapotranspiration; Eref) under different false positive rates for just the windows with non-repeated signatures (C) or for all windows (D).
